# Peripheral regeneration of Aβ low-threshold mechanoreceptors is limited despite activation of regenerative transcriptional pathways

**DOI:** 10.64898/2025.12.13.691790

**Authors:** Sara Bolívar, Natàlia Martínez-Mateu, David Ovelleiro, Esther Udina

**Author notes:** Corresponding author: Esther Udina.

## Abstract

Peripheral neurons can regenerate after nerve injury, yet functional recovery is often incomplete due to non-specific or incomplete target reinnervation. Among sensory neuron subtypes, Aβ low-threshold mechanoreceptors (Aβ-LTMRs) mediate discriminative touch, a modality often incompletely restored after nerve injury. However, the mechanisms governing Aβ-LTMR regeneration remain largely unexplored. Here, we characterised the anatomical and transcriptional features of Aβ-LTMRs regeneration after nerve injury. We assessed the extent of axonal regeneration after sciatic nerve crush and the preferential regeneration following femoral nerve transection using Calb1-Cre/tdTomato reporter mice. In both models, regeneration was incomplete, failing to reach control values. Moreover, Aβ-LTMRs preferentially reinnervated cutaneous rather than muscle pathways. To define the molecular program underlying this response, we performed ribosome-bound RNA sequencing from Calb1-Cre/Ribotag mice 7 days after nerve crush. Aβ-LTMRs upregulated canonical regeneration-associated genes, including *Atf3, Sprr1a, Gal*, and *Gap43*. Comparison with published RNA-seq datasets from other sensory neuron populations revealed only modest overlap, indicating that the core injury response shared between neurons is accompanied by a neuron subtype-specific programme. Together, these findings define the regenerative profile of Aβ-LTMRs across anatomical and transcriptomic levels, revealing the limited regenerative capacity of this neuron subpopulation despite robust activation of classical regeneration-associated genes.

## INTRODUCTION

Nerve injuries result in the loss of motor and sensory functions in the region that was innervated by the damaged nerve. Despite the capacity of peripheral axons to regenerate, functional recovery usually remains incomplete. The slow regeneration rate can delay the recovery and, even if axons reach distal targets, the lack of specificity in reinnervation can impair the function chronically. To achieve a successful regeneration, motor axons should reinnervate muscles, while sensory neurons need to reach sensory targets. Furthermore, sensory neurons in the dorsal root ganglia (DRG) are heterogeneous, having different targets, functions and transcriptional profiles (Usoskin et al., 2015; Qi et al., 2024). Given these considerations, understanding the mechanisms governing regeneration and axon guidance in specific cell-types is necessary to improve the functional recovery after a nerve injury.

Low-threshold mechanoreceptors (LTMRs) are a subtype of primary afferents that are activated by innocuous mechanical forces, encoding for light touch, pressure and vibration sensation. Not much is known about the regeneration of this subpopulation, partially due to the lack of good labelling strategies to study these neurons. Classical studies have primarily distinguished between myelinated and unmyelinated fibres, suggesting that small neurons exhibit superior regenerative capacity (Krarup et al., 1989; Navarro et al., 1994; Lozeron et al., 2004). Understanding the behaviour of A-LTMRs after injury can be crucial to recover fine sensory functions, such as spatial discrimination and fine motor interactions. Furthermore, A-LTMRs can induce mechanical allodynia after nerve injury (Xu et al., 2015; Dhandapani et al., 2018; Gautam et al., 2024), and their lack of regeneration can lead to aberrant C-fibre reinnervation that causes neuropathic pain (Gangadharan et al., 2022; Jeon et al., 2024). Thus, promoting accurate regeneration of A-LTMRs could be crucial to both improve functional recovery and reduce neuropathic pain after nerve injury.

Over the last decade, efforts have been made to identify specific markers and genetic tools for studying peripheral neuron sub-populations. Single-cell (sc) and single-nucleus RNA sequencing (snRNA-seq) studies have identified at least 15 transcriptionally defined neuronal clusters in the DRG (Usoskin et al., 2015; Zeisel et al., 2018; Zheng et al., 2019; Renthal et al., 2020; Zhang et al., 2022; Jung et al., 2023; Bhuiyan et al., 2024; Qi et al., 2024), highlighting the heterogeneity of primary afferents. From these markers, Calb1 emerged as a mouse genetic tool to label Aβ rapid adapting (RA)-LTMR (Zheng et al., 2019), which opens an opportunity for investigating the physiopathology of regeneration in these neurons.

In our previous work, we made a comparative characterisation of the regeneration of different neurons subtypes after nerve injury, including motoneurons, proprioceptors and nociceptors (Bolívar et al., 2024). We found a significant difference in the regeneration rate of these populations: proprioceptors showed the slowest regeneration rate, whereas nociceptors regenerated quicker. We found these functional differences were accompanied by distinct transcriptional profiles after nerve injury that could potentially explain the variable regeneration between neuron subtypes. Here, we aimed to extend our prior analysis by characterising the regeneration profile of an Aβ RA-LTMRs sub-population. We used the genetically labelled mouse line Calb1-Cre/tdTomato (tdT) to define the regeneration rate and preferential reinnervation of these neurons after nerve injury. We then took advantage of the RiboTag mouse to isolate ribosome-bound transcripts from this neuronal subpopulation (Sanz et al., 2009, 2019) and examine their transcriptional response to injury. Our results demonstrate that Aβ RA-LTMRs exhibit limited regeneration *in vivo* following sciatic crush and femoral transection injuries. We further show that regenerating axons preferentially regenerate towards cutaneous pathways after transection. Notably, despite their modest regenerative capacity, this subpopulation activates a transcriptional program associated with regeneration after injury, including significant upregulation of regeneration-associated genes (RAGs).

## METHODS

### Animals

Double transgenic mice were generated by breeding homozygous Calb1-2A-dgCre-D (#023531, The Jackson Laboratory, RRID:IMSR_JAX:023531) with homozygous Ai9(RCL-tdT) (#007909, The Jackson Laboratory, RRID:IMSR_JAX:007909) or heterozygous Ribotag mice, kindly donated by Dr. Elisenda Sanz (Universitat Autònoma de Barcelona, Spain). Cre expression was induced by intraperitoneal injection of trimethoprim (50 mg/ml; Sigma #T78883) dissolved in DMSO. Mice were treated for 3 consecutive days with a dose of 100 μg per gram of body weight per day and used at least 48 hours (Calb1^Cre^/tdT) or 2 weeks (Calb1^Cre^/Ribotag) after the injections.

Mice were housed in a controlled environment (12-hour light-dark cycle, 22±2ºC), in open cages with water and food *ad libitum*. All experimental procedures were approved by the Universitat Autònoma de Barcelona Animal Experimentation Ethical Committee and followed the European Communities Council Directive 2010/63/EU and the Spanish National law (RD 53/2013).

### Immunofluorescence

Adult mice (8-to 12-weeks old) were euthanized with intraperitoneal sodium pentobarbital (30 mg/kg) and perfused with 4% PFA in phosphate buffered saline (PBS). Lumbar DRGs, footpads, and hairy skin were removed and stored in PBS containing 30% sucrose at 4ºC for later processing. DRGs were serially cut in a cryostat (15 μm thick) and picked up on glass slides, whereas footpads were cut (50 μm thick) and stored free-floating. Samples were hydrated with PBS and permeabilized with PBS with 0.3% triton. Sections were blocked with 10% normal donkey serum in PBST for 1 hour at room temperature and then incubated with the primary antibody in PBST overnight at 4ºC (Table 1). Sections were washed with PBST and further incubated with a specific secondary antibody bound to Alexa 488 (1:200, Invitrogen) for 2 hours at room temperature (DRGs) or overnight at 4ºC (skin). For IB4 immunostaining, an overnight incubation at 4ºC with anti-lectin I was done prior to the secondary antibody incubation. Finally, after washing in PBS, slides were mounted with Fluoromount-G mounting medium (Southern Biotech, 0100-01) and imaged with a confocal microscope (Leica SP5) or an epifluorescence microscope (Olympus BX51, Olympus, Hamburg, Germany) equipped with a digital camera (Olympus DP50, Olympus, Hamburg, Germany). We counted the number of neurons tdTomato^+^ and how many of them co-labelled with the different markers using ImageJ software. Small (diameter <25 μm) and medium-to-large (diameter ≥25 μm) neurons were quantified separately. For size distribution, we contoured 100 neurons using ImageJ software.

**Table 1.**

| Sample | Thickness | Immunofluorescence |
| --- | --- | --- |
| DRG | 15 μm | Parvalbumin (1:1000; Swant Cat# PV-28, RRID:AB_2315235) |
|  |  | Neurofilament H (1:1000; Biolegend Cat# 801701, RRID:AB_2564642) |
|  |  | CGRP (1:200; Sigma-Aldrich Cat# PC205L, RRID:AB_2068524) |
|  |  | Calbindin D-28K (1:200, Millipore Cat# AB1778, RRID:AB_2068336) |
|  |  | Tyrosine hydroxylase (1:250; Millipore Cat# AB1542, RRID:AB_90755) |
| | | Isolectin B4 (10 μg/ml; Vector Cat# L-1104, RRID:AB_2336498) |
|  |  | Anti-lectin I (1:500; Vector Cat# AS2104, RRID:AB_2314660) |
| Skin | 50 μm | PGP 9.5 (1:500; Springbio Cat# E3340, RRID:AB_1661545) |

### Femoral nerve transection and retrograde tracing

Adult Calb1-Cre/tdT mice (6 males and 11 females, 8-to 12-weeks old) were anesthetized with intraperitoneal ketamine (90 mg/kg) and xylazine (10 mg/kg) and the left femoral nerve was exposed using an inguinal approach. The subcutaneous adipose tissue was carefully removed until the nerve and its ramifications were accessible. Then, the femoral nerve was cut above the bifurcation, 6 mm proximal to the entry of the quadriceps branch into the quadriceps muscle. Immediately afterward, the injured nerve was repaired with 5-10 μl of fibrin glue. The glue was prepared by mixing thrombin (25 U/ml, ICN Biomedicals, Cat# 0215416305) in calcium chloride (45 mmol/L, Sigma, Cat# C1016), human fibrinogen (100 mg/ml, Sigma, Cat# F3879), and bovine fibronectin (8 mg/ml, Sigma, Cat# F4759) in a 2:1:1 ratio (Guest et al., 1997; Akhter et al., 2019). When the fibrin glue polymerized, the skin was closed with a 6-0 nylon suture (Aragó). Animals were monitored periodically until the end of the experiments.

After 1 or 8 weeks, mice were anesthetized to re-expose the left femoral nerve (1 week: n=6, 8 weeks: n=8). Additionally, some uninjured mice were included in the process as controls (n=3). The quadriceps and the saphenous branches were cut at the same distance from the injury (approximately at 6 mm), just before the entry of the quadriceps branch into the muscle. The end of the muscle and cutaneous branch was soaked randomly in either Fluorogold (4%, Fluorochrome, Cat# 52-9400) or True Blue Chloride (2.5%, Setareh Biotech, Cat# 7120), and protected from the light for 45 minutes. To avoid spilling, we set up a small well using a piece of parafilm covered by a Vaseline circumference, we introduced the nerve stump into the well, and we filled it with the retrograde tracers.

One week later, mice were euthanized with intraperitoneal pentobarbital (30 mg/kg) and perfused with 4% PFA in PBS. The ipsilateral L3 and L4 DRGs were extracted and stored in PBS with 30% sucrose at 4ºC until they were serially cut in a cryostat (Leica, 15 μm thick) in and picked up in glass slides. We imaged DRGs with an epifluorescence microscope (Olympus BX51, Olympus, Hamburg, Germany) equipped with a digital camera (Olympus DP50, Olympus, Hamburg, Germany) and CellSens Digital Imaging software (version 1.9, Olympus, Hamburg, Germany). We counted the number of Calb1^+^ neurons that regenerated using the muscle or the cutaneous branch by counting the tdTomato^+^/TrueBlue^+^ or tdTomato^+^/Fluorogold^+^ neurons in the DRG. The experimenter was blinded to the tracer-branch sequence for each animal.

### Sciatic nerve crush

Adult Calb1-Cre/tdT mice (3 males and 11 females, 8-to 12-weeks old) were anesthetized with intraperitoneal ketamine (90 mg/kg) and xylazine (10 mg/kg) and the right sciatic nerve was exposed through a gluteal muscle-splitting incision. Nerves were crushed 3 mm distal to the sciatic notch with fine forceps (Dumont no. 5) applied for 30 seconds. The lesion site was labelled with an epineural suture stitch (10-0 nylon suture, Alcon) and the muscle and the skin were closed (6-0 nylon suture, Aragó). Animals were monitored periodically until the end of the experiments. After 7 or 9 days, mice were euthanized with intraperitoneal pentobarbital (30 mg/kg) and perfused with 4% PFA in PBS (n=7 for each day). The right sciatic nerve and its extension as tibial nerve were harvested and stored in PBS with 30% sucrose. We also collected some contralateral nerves as control. A segment from 12 mm to 17 mm from the lesion site was serially cut in 10 μm longitudinal sections in a cryostat (Leica) and picked up in glass slides. The slides were mounted with Fluoromount-G mounting medium (SouthernBiotech, #0100-01) and visualized in an epifluorescence microscope (Nikon Eclipse Ni-E, Nikon, Tokyo, Japan) equipped with a digital camera (Nikon DS-RiE, Nikon, Tokyo, Japan) and Nikon NIS-Element BR software (version 5.11.03, Nikon, Tokyo, Japan). Using the software, we drew a line perpendicular to the nerve and counted the number of axons that crossed the line at 12 and 17 mm from the injury in one out of three sections.

### Translating ribosome affinity purification

Adult Calb1-Cre/Ribotag mice (total of 12 females and 9 males, 8 to 11 weeks old) were anesthetized with intraperitoneal ketamine (90 mg/kg) and xylazine (10 mg/kg), and the right sciatic nerve was crushed as detailed above. Then, the femoral nerve was exposed and crushed for 30 s above the bifurcation. The skin was closed with a 6-0 nylon suture (Aragó) and animals were monitored periodically until the end of the experiments. After 7 days, mice were euthanized with intraperitoneal pentobarbital (30 mg/kg), and L3-L5 DRGs were rapidly dissected. Samples were placed in cold Gey’s solution enriched with glucose (6 mg/ml) until they were used for the Ribotag assay. L3-L5 DRGs from groups of 3 mice were pooled and homogenized in 1 ml of homogenization buffer as described previously (Sanz et al., 2009). Female and male mice were used for the study, pooled in groups of the same sex. After centrifugation of the homogenate, 40 μl of the supernatant was stored as an input sample, whereas 4 μl of anti-HA antibody (Covance, Cat# MMS-101R) was added to the remaining lysate and incubated for 4 hours at 4ºC with rotation. Then, 200 μl of protein A/G magnetic beads (ThermoFisher, Cat# 88803) were washed and added to the lysate for 5 hours at 4ºC with rotation. Samples were washed in a high-salt buffer to weaken the antibody-magnetic bead binding and beads were removed using a magnet. RNA was isolated from the samples using the RNeasy Micro Kit (Qiagen, Cat# 74004) and quantified with Quant-it RiboGreen RNA Assay Kit (ThermoFisher, Cat# R11490). The integrity of the RNA was assessed using the 2100 Bioanalyzer system with the RNA 6000 Nano or Pico chips (Agilent Technologies).

### Library preparation and RNA-sequencing

#### Library preparation and RNA sequencing

Lexogen next-generation sequencing (NGS) services (Vienna, Austria) were used to perform RNA quality control and RNA sequencing. Libraries were prepared via the CORALL mRNA-Seq V2 preparation kit. All the samples were sequenced via the Illumina NextSeq 500 platform in runs of 2×150 bp, yielding at least 15 million reads per sample. FASTQ files were evaluated for quality via FastQC (v0.11.9). Reads were aligned to the coding DNA reference database (Ensembl Mouse database, Genome assembly: GRCm39, release 103) using Salmon (v1.4.0). A total of 64 single-end samples were quantified. The quantified transcript reads were mapped to genes and imported to the R environment (version 4.4.0) using the library “tximport” (v 1.32.0). Then, the “DESeq2” library (v1.44.0) was used to perform the differential expression analysis. Volcano plots were plotted using the R library “EnhancedVolcano” (v 1.22.0). Only genes with more than 100 counts among all samples were used in the differential analysis.

Gene set enrichment was performed using the R library “STRINGdb”5, using the databases offered by the String platform6. Only proteins with a P-value lower than 0.05 are used in the enrichment. Pathway analysis was made using Bioconductor libraries “KEGGREST” and “pathview”. For each groups comparison a Wilcoxon test is performed for each pathway, producing a list of KEGG pathways with an associated p-value. The calculation of this p-value takes into account the original p-value and fold change for a given gene and the number of genes in each KEGG pathway. For Gene Set Enrichment Analysis (GSEA) we customized a list of RAGs that have been described extensively in the literature (van Kesteren et al., 2011; Allodi et al., 2012; Ma and Willis, 2015) to evaluate if these genes were differentially and concordantly expressed seven days after injury. The list of RAGs was: *Adcyap1, Arg1, Atf3, B2m, Basp1, Bdnf, Bid, Ccl2, Cdc42, Cdkn1a, Cebpd, Calca, Creb1, Dpysl2, Drd4, Fmr1, Gal, Gap43, Gdnf, Hspb1, Il6, Itgb5, Itgb7, Jun, Klf7, Lgals1, Mtl5, Ngfr, Rac1, Smad1, Snip1, Sox11, Stat3, Syn1, Timp1, Trp53*. GSEA was performed by using the R Package Cluster profiler from BiocManager.

Correlations with our previous RNA-seq dataset were assessed using Pearson’s correlation. Genes with log2 fold-change values greater than 10 or less than –10 were excluded in this analysis to minimize potential confounding effects.

### Statistical analysis

GraphPad Prism 8 (version 8.0.2) was used for the statistical analysis of the histology data. The normal distribution of the samples was tested with Shapiro-Wilk test. The statistical test used in each analysis is specified in the results section. All data are expressed as group mean ± standard error of the mean (SEM). Differences were considered statistically significant if p<0.05.

## RESULTS

### Calb1-Cre/tdT mice preferentially label low-threshold mechanoreceptors

The Calb1-Cre transgenic mouse has been described as a tool for studying Aβ RA-LTMRs (Zheng et al., 2019). To corroborate this, we performed immunostaining in the DRG of Calb1-Cre/tdT reporter mice against markers of specific neuron subtypes (Figure 1). Surprisingly, we identified two clearly distinct fluorescent populations: medium-to-large neurons with a diameter above 25 μm (hereafter “large”) and small neurons below 25 μm (Figure 1b). Large neurons co-labelled predominantly with the Aβ-LTMR marker Calb1 (85.84 ± 2.85 %) and with NFH (85.58 ± 0.96 %). A high percentage of these neurons were also positive for the proprioceptive marker Parv (41.03 ± 4.51 %). In contrast, small neurons co-labelled mostly with Calb1 (89.59 ± 2.17 %), but only 6.54 ± 0.62 % were NFH^+^. These also did not show expression of Parv (1.57 ± 0.85 %). Neither large nor small neurons co-labelled with the nociceptive markers IB4 (0.29 ± 0.29 %) and CGRP (3.78 ± 2.05 %) (Figure 1c-e). These data are consistent with published scRNA-seq showing Calb1 expression in two distinct DRG neuron subtypes (Ntrk3^high^+Ntrk2 and Rxfp1 clusters, Supplementary figure 1).

**Figure 1.**
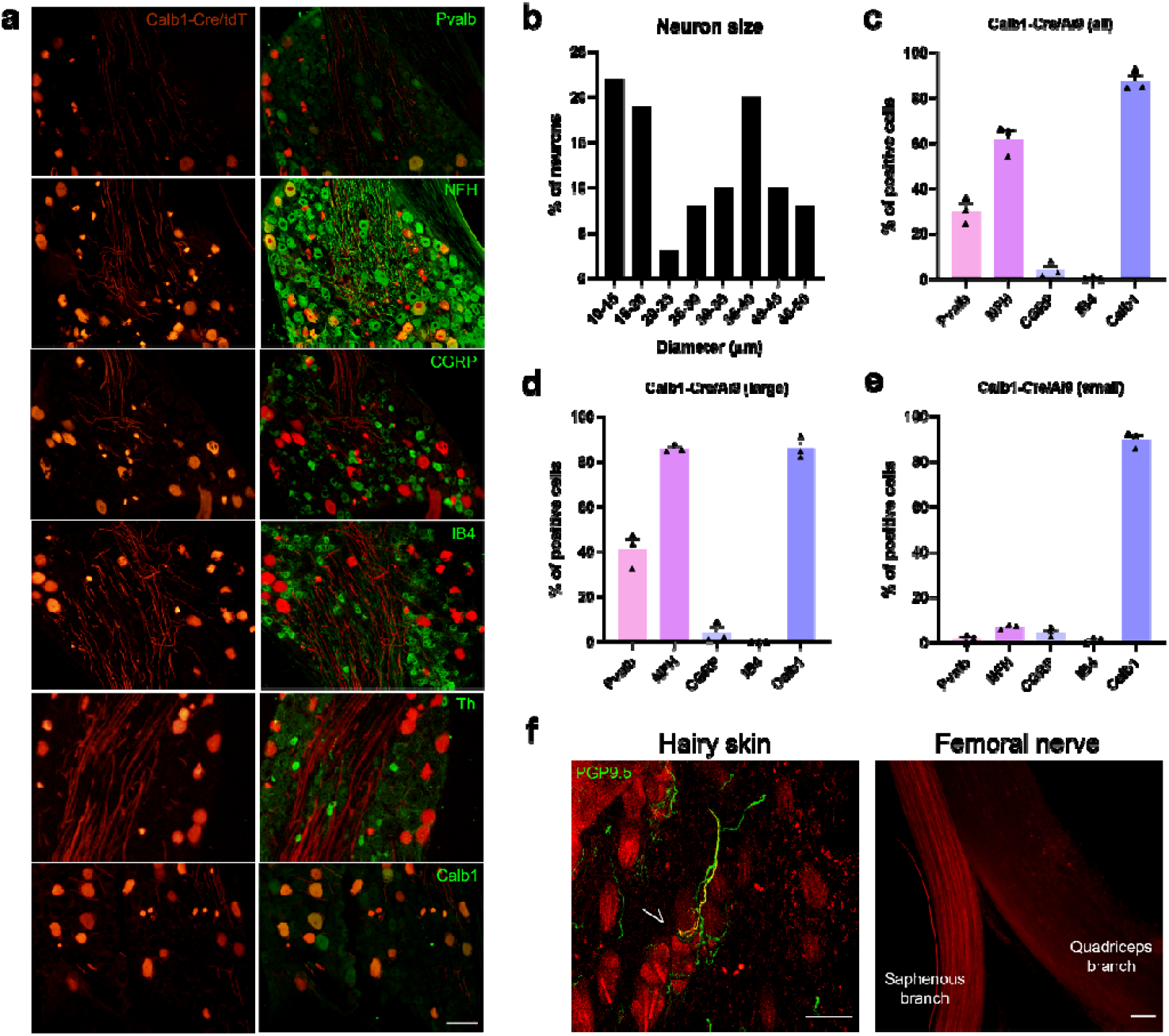
Characterization of Calb1-Cre/tdT mice. a) Immunostainings against specific markers in the DRG (parvalbumin, Pvalb; neurofilament heavy chain, NFH; calcitonin gene-related peptide, CGRP; isolectin B4, IB4; tyrosine hydroxylase, Th; calbindin-1, Calb1). b) Size distribution of tdTomato^+^ neurons in the DRG. c-e) Quantification of tdTomato co-localization with the indicated markers in c) all neurons, d) large neurons (>25 μm diameter) and e) small neurons (<25 μm) (n=3). f) Representative images of hairy skin and femoral nerve. The arrow indicates a tdTomato^+^ axon (yellow) associated with a hair follicle (PGP9.5 immunostaining is shown in green). Scale bar: 100 μm.

Next, we evaluated axonal distribution in the femoral nerve and skin. We found that most fluorescent axons in the femoral nerve were in the saphenous branch. In hairy skin, axons were found in hair follicle receptors (Figure 1f). Altogether, these observations indicate that fluorescent neurons in Calb1-Cre/tdT mice fit the profile of cutaneous afferents, labelling predominantly (although not exclusively) Aβ RA-LTMRs.

### Aβ-LTMR preferentially regenerate toward cutaneous targets

We assessed whether Calb1^+^ neurons show preferential regeneration into a specific nerve branch after injury. We transected and repaired the femoral nerve above its bifurcation and, using retrograde tracers, quantified neurons that had regenerated through the muscle branch, the cutaneous branch, or both at 1 and 8 weeks post-injury (Figure 2a,b). In intact nerves, we corroborated that most retrotraced neurons projected to the cutaneous branch (83.15 ± 4.69 %), as expected for a population enriched in Aβ-LTMRs. One week after injury, we did not observe significant differences between the number of neurons found in the muscle and the cutaneous branch (14.17 ± 3.37 neurons and 22.33 ± 5.32 neurons, respectively). Conversely, by 8 weeks neurons showed a preferential regeneration toward the cutaneous branch (25.13 ± 2.42 neurons in muscle branch vs 39.5 ± 5.43 neurons in the cutaneous branch, p=0.039) (Figure 2c). Although modest (59.96 ± 4.32 % of neurons in the cutaneous branch; Figure 2d), this preference was consistent across most animals (Table 2). Thus, Aβ-LTMRs show a preferential regeneration towards their original branch comparable to that described for motoneurons and other sensory afferents.

**Table 2.**
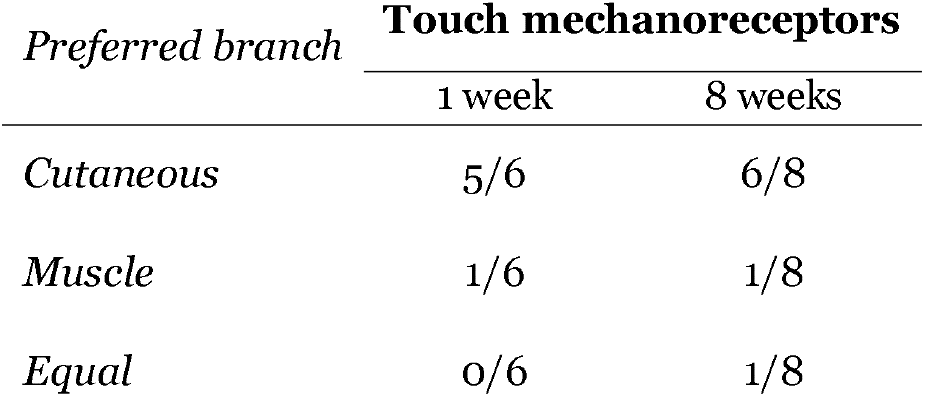

**Figure 2.**
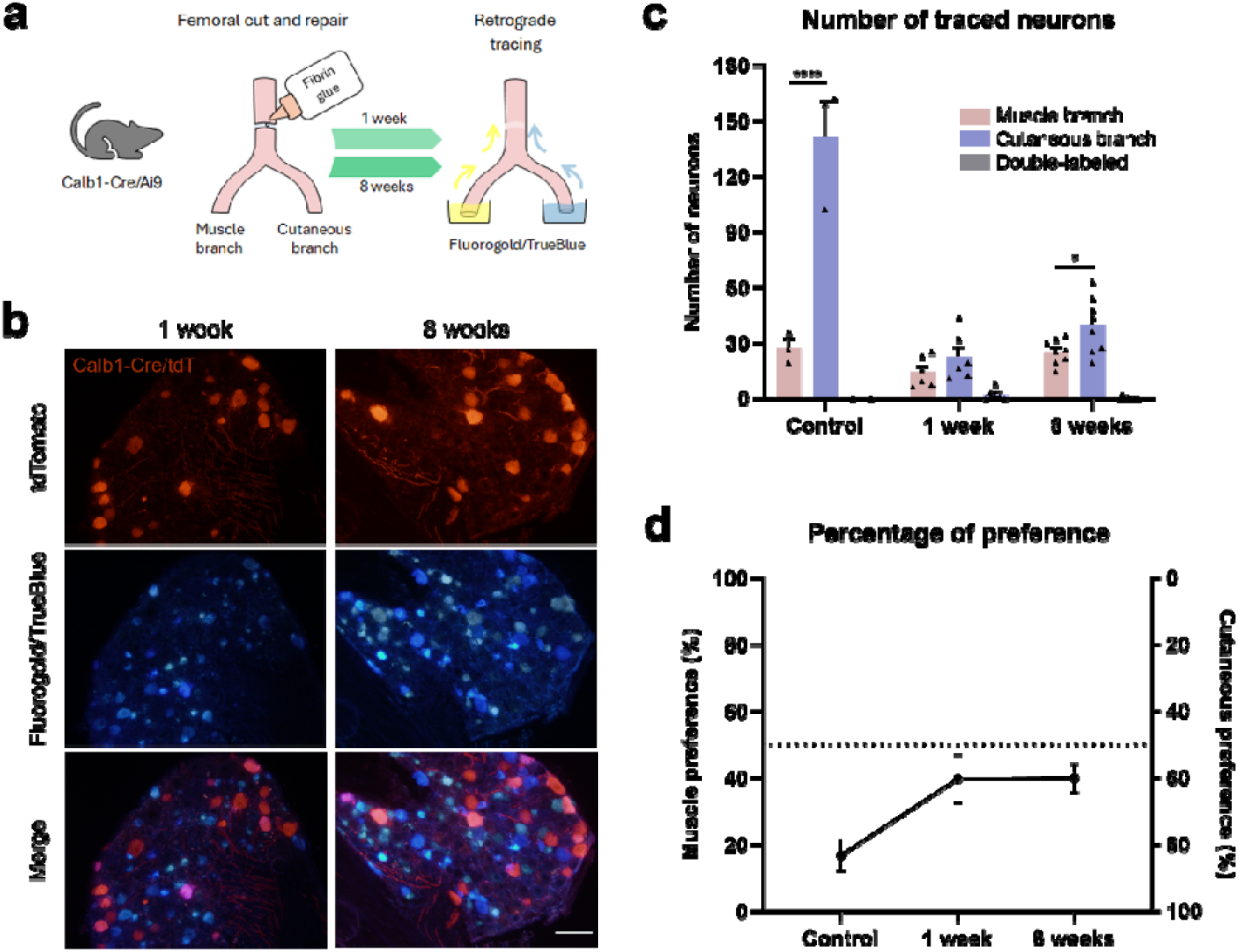
Preferential regeneration of Calb1^+^ neurons. a) Schematic representation of the experimental design. b) Representative DRG images at 1 week (n=6) and 8 weeks (n=8) after nerve transection. Neurons were counted when tdTomato (red) co-localized with TrueBlue (blue) or Fluorogold (yellow). c) Quantification of neurons whose axons projected to the muscle branch, the cutaneous branch, or both. d) Percentage of retrotraced neurons in the muscle or cutaneous branch calculated over the total number of regenerated neurons. Muscle preference is plotted on the left y-axis and cutaneous preference on the right y-axis. Differences between branches were assessed by two-way ANOVA followed by a Sidak post hoc test. *p<0.05, ****p<0.0001 (muscle vs cutaneous branch). Scale bar: 100 μm.

### Calb1^+^ neurons have limited regeneration after nerve injury

To establish the regenerative capacity of Calb1^+^ neurons, we performed a sciatic nerve crush and quantified the number of tdTomato^+^ axons at two distances distal from the lesion site (Figure 3a). At 7 days post-injury (dpi), only 29.61 (± 5.24) % and 5.98 (± 1.65) % of the total axons had reached 12 and 17 mm, respectively. By 9dpi, regeneration increased to 59.47 (± 9.13) % at 12 mm and 51.60 (± 6.96) % at 17 mm (Figure 3c-d). At both distances, there was a significant increase in the number of axons between 7 dpi and 9 dpi (p=0.003 at 12 mm and p<0.001 at 17 mm), but none of the conditions reached control values. Notably, at 9 dpi there was no significant difference between 12 mm and 17 mm (p = 0.706), consistent with a regeneration plateau. Incomplete regeneration was also evident after femoral nerve transection and repair, where only 23.02 ± 4.14% and 38.69 ± 3.52% of neurons regenerated into distal branches at 1 and 8 weeks, respectively (Figure 3e). Together, these data indicate that Aβ-LTMRs display limited regenerative capacity and fail to fully recover after peripheral nerve injury.

**Figure 3.**
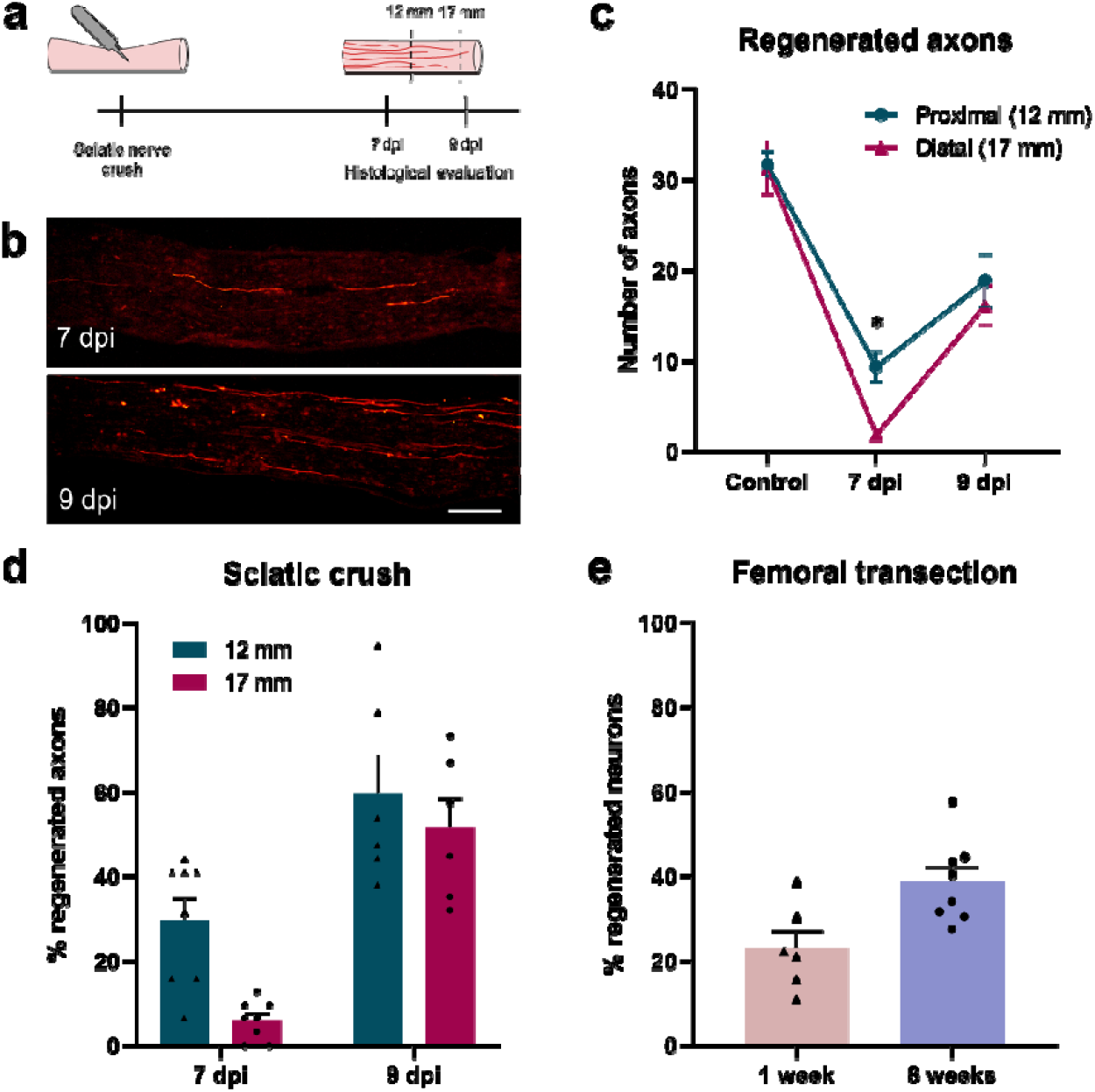
Regenerative capacity of Aβ-LTMR. a) Schematic representation of the experimental design. b) Longitudinal sections of tibial nerves from Calb1-Cre/tdT mice at 7 days (n=8) and 9 days (n=6) post-injury (dpi). c) Number and d) percentage of tdTomato^+^ axons in the tibial nerve after injury. e) Percentage of neurons that regenerated 1 or 8 weeks after femoral transection, assessed by retrograde tracers. Percentages were normalized to uninjured controls. Differences in axon regeneration were assessed by two-way ANOVA followed by a Sidak post hoc test. *p<0.05 (proximal vs distal). Scale bar: 100 μm.

### Regenerative transcriptional program activated by Aβ-LTMR

We isolated the pool of actively translated mRNAs in Calb1^+^ neurons using the Ribotag assay (Fig 4a). Calb1-Cre mice were crossed with RiboTag mice to generate double-transgenic animals expressing a hemagglutinin (HA) tag specifically in ribosomes of this neuronal population. By immunoprecipitating the tagged ribosomes from DRG homogenates, we isolated neuron-specific mRNAs and performed RNA-seq to compare naïve vs nerve crush samples.

**Figure 4.**
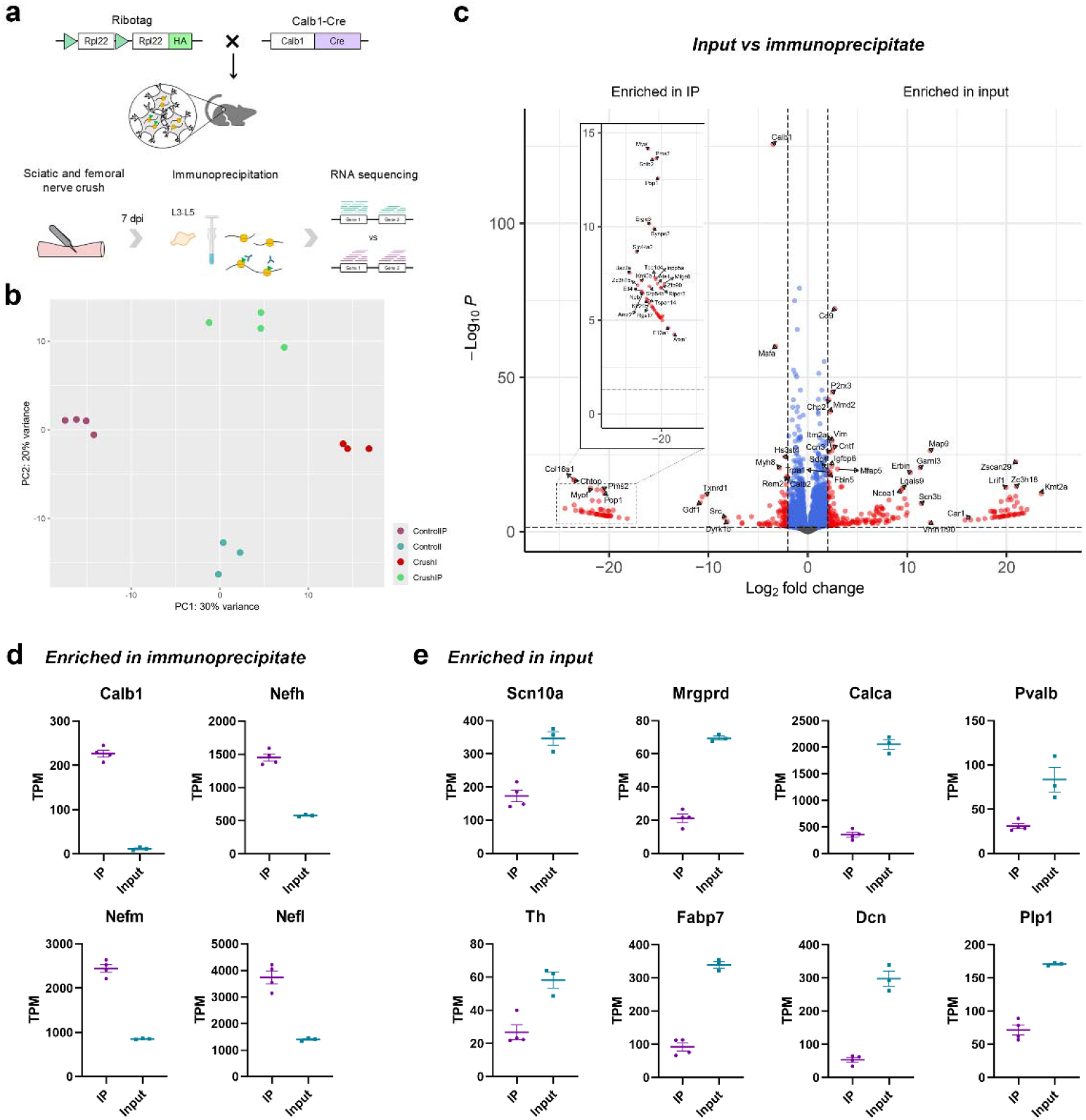
Validation of the immunoprecipitate specificity in naive Calb1-Cre/Ribotag mice. a) Schematic representation of the experimental design. b) Principal component (PC) analysis of the samples showing segregation of samples by experimental group. c) Volcano plot comparing input (whole DRG) versus immunoprecipitate (IP; Calb1^+^ neurons) showing significant differentially expressed genes in blue (log_2_ fold change <2) and red (log_2_ fold change >2). d) Transcripts per million (TPM) for representative genes enriched in the IP or e) enriched in the input.

To assess the specificity of the strategy, we confirmed that the RNA from control immunoprecipitates (IPs) had enriched transcripts of the well-established Aβ-LTMRs transcripts *Calb1, Nefh, Nefm* and *Nefl* compared to the input (whole DRG) (Figure 4d). Conversely, IPs were depleted for markers of other neuronal subtypes (*Scn10a, Mrgprd, Calca, Pvalb, Th*), as for glial markers (*Fabp7, Dcn, Plp1*) (Figure 4e). Principal component analysis (PCA) revealed that samples clustered according to their experimental group (Figure 4b).

Seven days after nerve crush, 1941 genes were differentially expressed in Calb1^+^ neurons (Figure 5a, Supplementary data 1). Despite the limited regeneration of these neurons, we found an enrichment of regeneration associated genes (RAGs) in the crush condition (*Sprr1a, Atf3, Gap43, Tubb2b*) (Figure 5B). Pathway analysis revealed significant activation of regeneration-related processes, including Axon guidance, Axonogenesis and Regulation of cell projection organization (Figure 5c,d).

**Figure 5.**
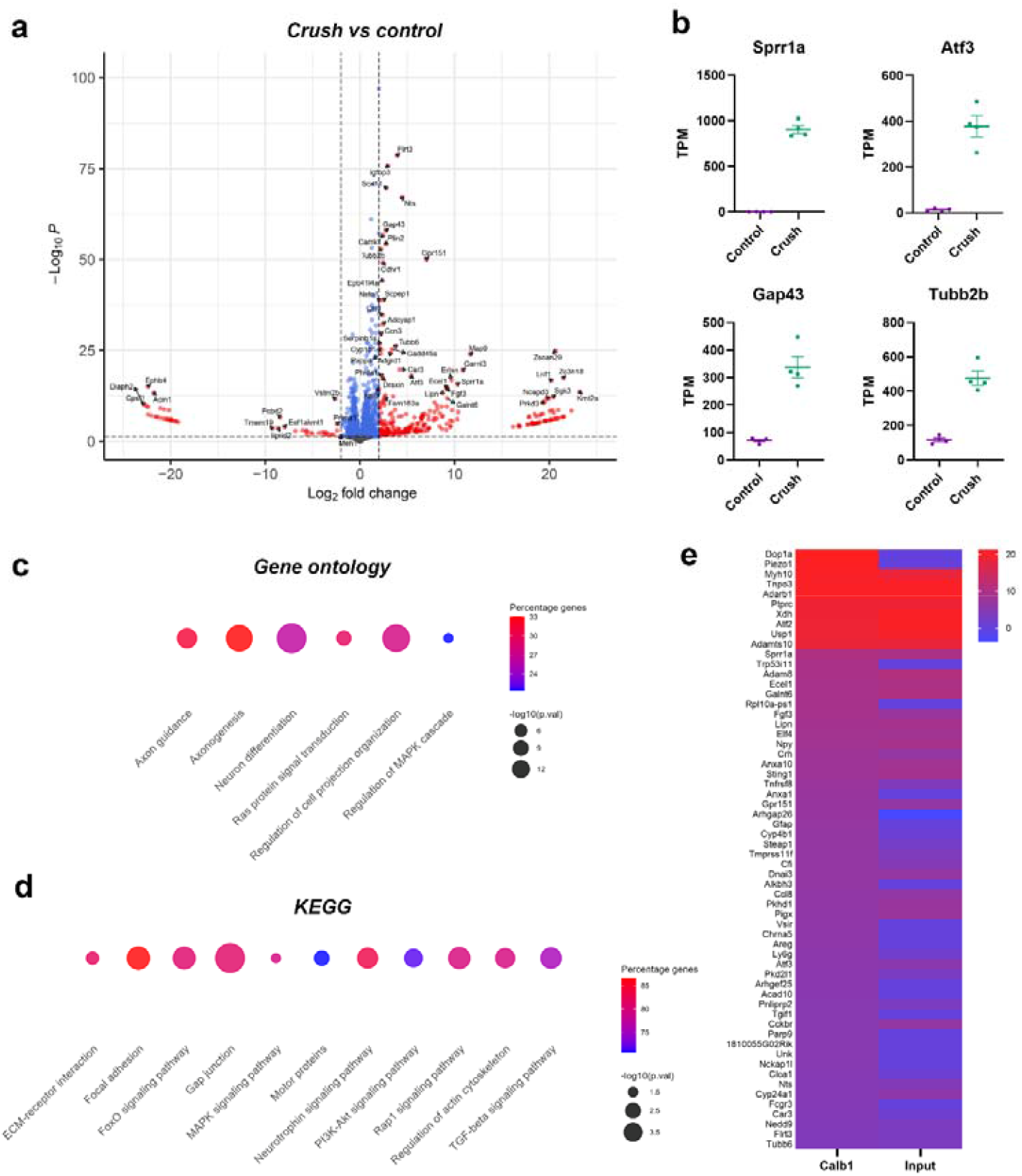
Injury-induced transcriptional changes in Aβ-LTMRs. a) Volcano plot of the control versus nerve crush DRG immunoprecipitates showing significantly differentially expressed genes in blue (log2 fold change <2) and red (log2 fold change >2). b) Transcripts per million (TPM) for representative genes enriched after crush. c) Gene ontology and d) Kyoto Encyclopedia of Genes and Genomes (KEGG) enrichment analysis showing pathways activated after nerve injury. e) Top upregulated genes in Calb1^+^ neurons after injury and their corresponding changes in input (whole DRG) samples (expressed in log_2_ fold-change).

Because peripheral neuron populations exhibit distinct regeneration behaviours despite sharing the same microenvironment, we hypothesized that population-specific transcriptional programs could underlie these physiological differences. Indeed, although some of the top upregulated genes after injury overlapped with those observed in the input, we found that many genes such as *Dop1a, Piezo1*, and *Trp53i11* displayed distinct activation patterns relative to input (Fig. 5e). This indicates that LTMRs engage a partially specific regenerative transcriptional program after nerve crush.

### Distinct transcriptional responses in peripheral neuronal subtypes

We have previously investigated the regeneration profile and transcriptional response to injury of different peripheral neuron populations using the same experimental approaches employed here for Aβ-LTMRs (Bolívar and Udina, 2022; Bolívar et al., 2024). Specifically, we analysed the transcriptional response of proprioceptors (Pvalb^+^ neurons), nociceptors (Trpv1^+^-lineage neurons), a population enriched for A high-threshold mechanoreceptors (Npy2r^+^-lineage) and motoneurons (MNs). We correlated the log_2_ fold change (log2FC) of differentially expressed genes after injury across populations and found a significant but modest correlation between gene expression in Calb1^+^ neurons and the other populations (Figure 6a-e), suggesting a partially distinct injury response. As expected, the Calb1 transcriptome correlated more strongly with the other sensory afferents than with motoneurons. Notably, the correlation was highest between Calb1^+^ and Pvalb^+^ neurons despite their distinct target organs, suggesting that similarities in function (low-threshold mechanosensation), size (medium to large neurons) and/or myelination may be more relevant to this response. Among the remaining populations, the strongest correlation was observed between Trpv1^+^ and Npy2r^+^, followed by Pvalb^+^ and motoneurons (Figure 6f).

**Figure 6.**
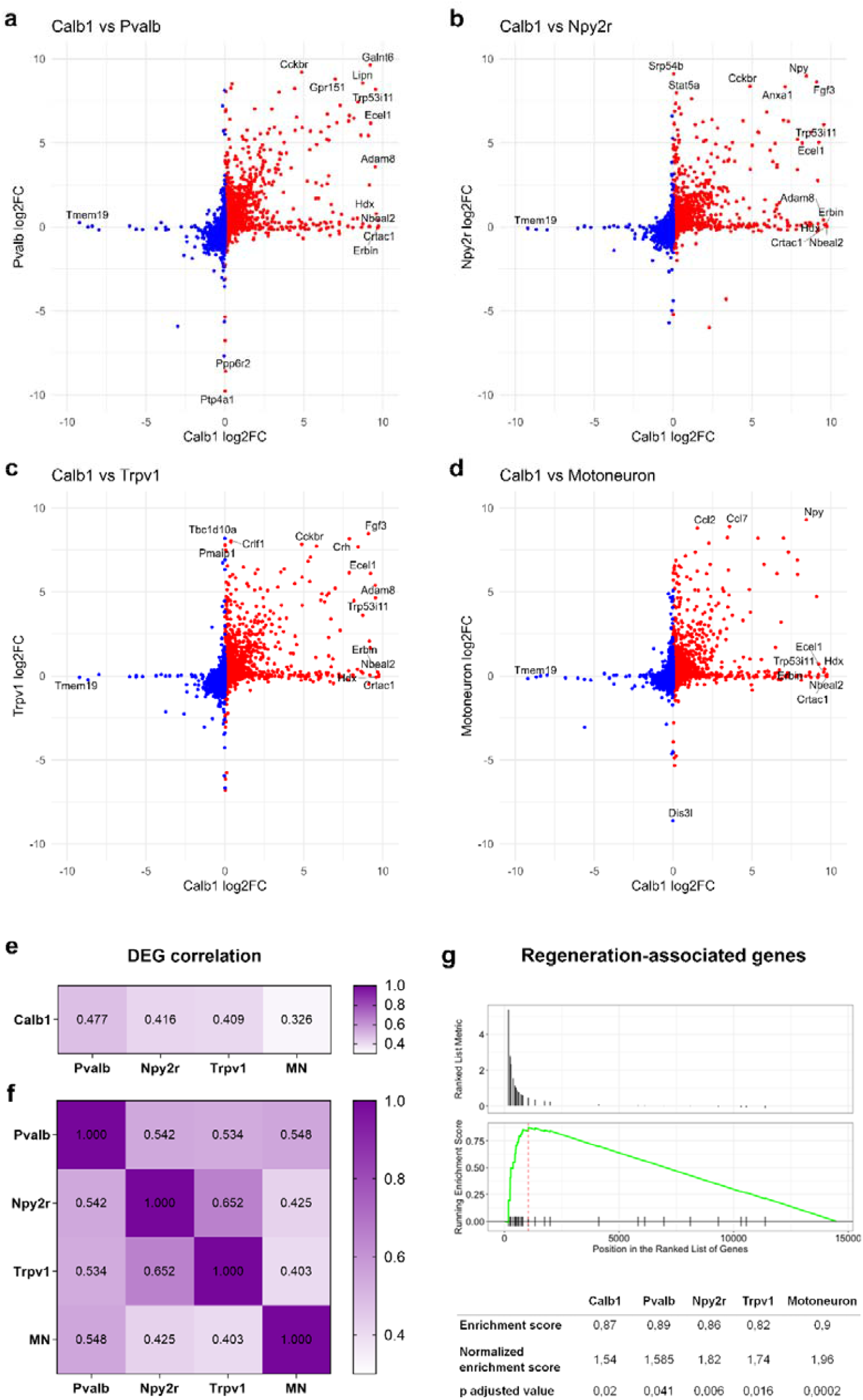
Cross-population comparison of injury-induced gene expression changes. a-d) Scatter plots comparing the log2foldchange (log2FC) of the differentially expressed genes (DEGs) after injury in Calb1+ neurons versus a) proprioceptors (Pvalb), b) an A high-threshold mechanoreceptor-enriched population (Npy2r), c) nociceptors (Trpv1) and d) motoneurons (MN). e) Heatmap of Pearson correlation coefficients (r) between DEGs in Calb1+ neurons and each neuron population. All correlations were significant (p<0.0001). f) Heatmap of pairwise Pearson correlations among neuronal subtypes from Bolívar et al. (2024). All correlations were significant (p<0.0001). g) Gene set enrichment analysis (GSEA) of a literature-curated set of regeneration-associated genes (RAGs) in Calb1+ neurons, with a summary of enrichment metrics across all populations.

Finally, we wondered whether the limited regeneration of LTMRs might reflect insufficient activation of RAGs. A targeted gene set enrichment analysis (GSEA) based on a literature-curated RAG list revealed significant positive enrichment of RAGs in Calb1^+^ neurons after injury (Figure 6g), comparable to that observed in the other populations examined (Supplementary Figure 2). Therefore, despite having a subtype-specific injury transcriptome, Aβ-LTMRs activate canonical RAGs to a similar extent across populations, indicating that RAG activation alone does not account for their limited regenerative capacity.

## DISCUSSION

The Calb1-Cre mouse line was originally described as a tool to label Aβ RA-LTMR (Zheng et al., 2019). In our hands, however, this line labelled two distinct neuronal subpopulations: one composed of large neurons and another of small neurons. We confirmed that large neurons exhibited expected features of Aβ RA-LTMR, including cutaneous targets and expression of histological markers associated with this neuron type. Interestingly, we found an additional labelled population of small-diameter, cutaneous neurons. Previous scRNA-seq studies have identified a Rxfp1^+^ cluster expressing Calb1. These Rxfp1^+^ neurons represent a poorly characterised, low-abundance sensory neuron subtype (Zeisel et al., 2018; Wang et al., 2021, 2023). The discrepancy with previous reports may result from differences in the timing of trimethoprim-induced transgene activation, as previous studies induced expression at P18–P19 (Zheng et al., 2019), whereas our induction was performed in adult mice (7–9 weeks old). As our main goal was to improve target selectivity, we analysed both populations together as cutaneous neurons. Future studies could leverage recently developed genetic tools (Qi et al., 2024) to investigate more precisely defined subtypes.

After a nerve transection, endoneurial tubes are disrupted and regenerating axons rarely re-enter their original pathways to the appropriate targets. However, it has been described that neurons can exhibit a preferential regeneration towards their original target organ (Brushart, 1988, 1993; Martini et al., 1994; Madison et al., 1996, 2007; Brushart et al., 1998; Robinson and Madison, 2003, 2005, 2006, 2013a, 2013b; Mears et al., 2003; Maki et al., 2005; Uschold et al., 2007; Bolívar and Udina, 2022; Li et al., 2025). Classically, this has been demonstrated in motoneurons, where axons preferentially grow through muscle pathways after transection. In our previous work, we confirmed that this phenomenon also occurs in some primary afferent subtypes, which show selectivity to regenerate towards their original target organ (Bolívar and Udina, 2022). Here we extended this work by showing that Aβ RA-LTMR preferentially regenerate into cutaneous pathways after injury. This preference varies across neuron subtypes: proprioceptors and motoneurons are attracted to muscle pathways, whereas cutaneous afferents prefer cutaneous branches. Several studies have described differences in the trophic factors secreted by denervated Schwann cells from muscle and cutaneous branches (Höke et al., 2006; Brushart et al., 2013), as well as molecular differences between Schwann cell subtypes (Martini et al., 1992, 1994; Saito et al., 2005, 2010; Jesuraj et al., 2012). These factors, together with trophic cues from target organs (Madison et al., 2007; Uschold et al., 2007), are likely key factors driving this regeneration selectivity. Nonetheless, all regenerating axons, regardless of subtype, are found in the same post-injury microenvironment and thus receive the same trophic support. Therefore, we believe that intrinsic differences in the regenerative programs activated by these subtypes should influence this process.

Similarly, we have consistently found that neuronal subtypes regenerate at different rates. Calb1^+^ neurons showed a strikingly low regeneration rate and even incomplete regeneration after a sciatic crush and femoral transection. This regeneration is comparable to that seen in proprioceptors and substantially poorer than that of nociceptors. Classical studies using cuff electrodes suggested that small afferents regenerate more effectively than large sensory neurons (Krarup et al., 1989; Negredo et al., 2004). Functional studies likewise indicated a faster, more complete functional recovery in unmyelinated compared to myelinated fibers (Navarro et al., 1994). More recently, incomplete regeneration of LTMRs after injury has been reported (Gangadharan et al., 2022; Jeon et al., 2024). This deficit is unlikely to be explained by neuronal loss, as large-diameter neurons are largely preserved after nerve injury (Cooper et al., 2024). This lack of regeneration can have detrimental consequences: beyond limiting functional recovery, insufficient regeneration may enable aberrant reinnervation of denervated territories by nociceptive fibers, contributing to neuropathic pain.

Does a differential transcriptional response to injury explain these differences? Previous snRNA-seq studies identified a conserved response across DRG neurons after nerve injury, with cells clustering into an “injured” transcriptional state (Renthal et al., 2020). Our data are consistent with the existence of a conserved core response, but they also reveal a subtype-specific regulation. Comparing the injury-induced translatome of Calb1+ neurons with our previous RNA-seq data from other populations we found only a modest correlation among DEGs. It is important to note that part of this divergence likely reflects baseline transcriptional differences between neuron subtypes. For example, injury-induced downregulation of subtype-identity markers will, by definition, be subtype specific. Yet many other DEGs could be key determinants of subtype-specific regeneration.

Given the limited regeneration capacity of Aβ-LTMR, we wondered whether this population shows reduced expression of classical RAGs. Instead, we observed a significant increase in RAGs, including Gap43, Sprr1a and Atf3. GSEA using a literature-based RAG list showed similar activation across all populations, including Trpv1-Cre/tdT which displays a much faster regeneration rate. Thus, transcriptional changes in canonical RAGs cannot explain differences in regeneration speed. We hypothesize that these differences may arise from differential regulation of additional, not-yet-characterized RAGs, or from divergence at the protein level (expression, localization, or post-translational modification) not captured by our approach. For example, Sting1 was selectively upregulated in Aβ-LTMR and proprioceptors after injury, the two subtypes with the poorest regeneration. Although the neuronal role of Sting1 is not completely understood, there is some evidence suggesting it can shape regeneration-degeneration dynamics. In experimental autoimmune encephalomyelitis, neuronal Sting1 activation has been linked to neurodegeneration in through the activation of ferroptosis (Woo et al., 2024), and ferroptosis was among the pathways enriched in our dataset after injury (Supplementary data 1). Furthermore, Sting1 global knockout has been shown to enhance functional recovery after nerve injury (Morozzi et al., 2021). These observations nominate Sting1 as a candidate modulator of regenerative capacity. In addition, we observed a subtype-specific upregulation of Atf2. Given prior evidence linking ATF2 to injury-evoked mechanical allodynia and thermal hyperalgesia (Salinas-Abarca et al., 2018), this gene may be relevant to subtype-specific consequences of injury.

Alternatively, size and the degree of myelination may explain some of the injury-specific responses. During Wallerian degeneration, myelin debris is cleared by macrophages and Schwann cells. Inhibitory myelin-associated molecules and extracellular matrix components, such as myelin-associated glycoprotein or chondroitin sulphate proteoglycans, can hinder axon regeneration (Mukhopadhyay et al., 1994; Torigoe and Lundborg, 1998; Niederöst et al., 2002; Zuo et al., 2002; Mears et al., 2003; Sarhane et al., 2019). In this context, unmyelinated axons might have a physical advantage, as their endoneurial tubes require less debris clearance. Together with the greater energetic demands of larger, myelinated axons, this could contribute to slower regeneration of myelinated sensory afferents.

This work has some limitations. First, we used Calb1-Cre mice to study Aβ-LTMRs. Although this line preferentially labels Aβ-LTMRs, it also labels a second cutaneous population. While this data is relevant for target selectivity, more refined genetic tools would help isolate Aβ-LTMRs with greater specificity. Second, we analysed the transcriptome at a single time point (7 days after sciatic crush). A time course and additional injury paradigms (e.g. nerve transection) would better define the dynamics and regeneration profile of the Aβ-LTMR response. Finally, examining the transcriptome at the axonal compartment would be valuable, as local translation at growth cones may be particularly relevant for the regenerative response and could help explain subtype-specific differences in regeneration after nerve injury.

## Acknowledgements

The authors appreciate the technical help of Neus Hernández. Mònica Espejo and Jessica Jaramillo.

