## Supplementary figures for "Peripheral regeneration of Aβ low-threshold mechanoreceptors is limited despite activation of regenerative transcriptional pathways"

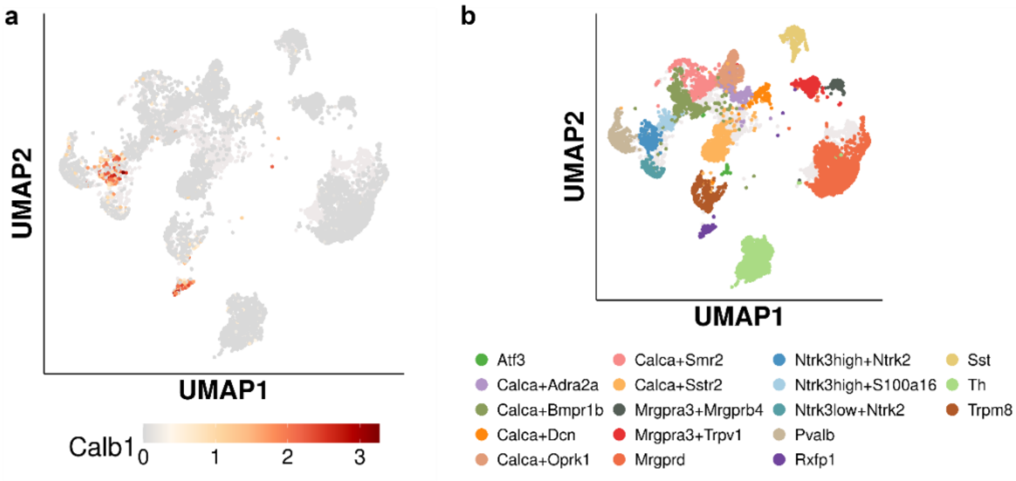


**Supplementary figure 1. Calb1 gene expression in DRG neurons from Bhuiyan et al. (2024) scRNA-seq data.** a) Calb1 expression is restricted to two distinct neuronal clusters. b) Cluster annotations identify these Calb1$^{+}$ clusters as Ntrk3high+Ntrk2 and Rxfp1.

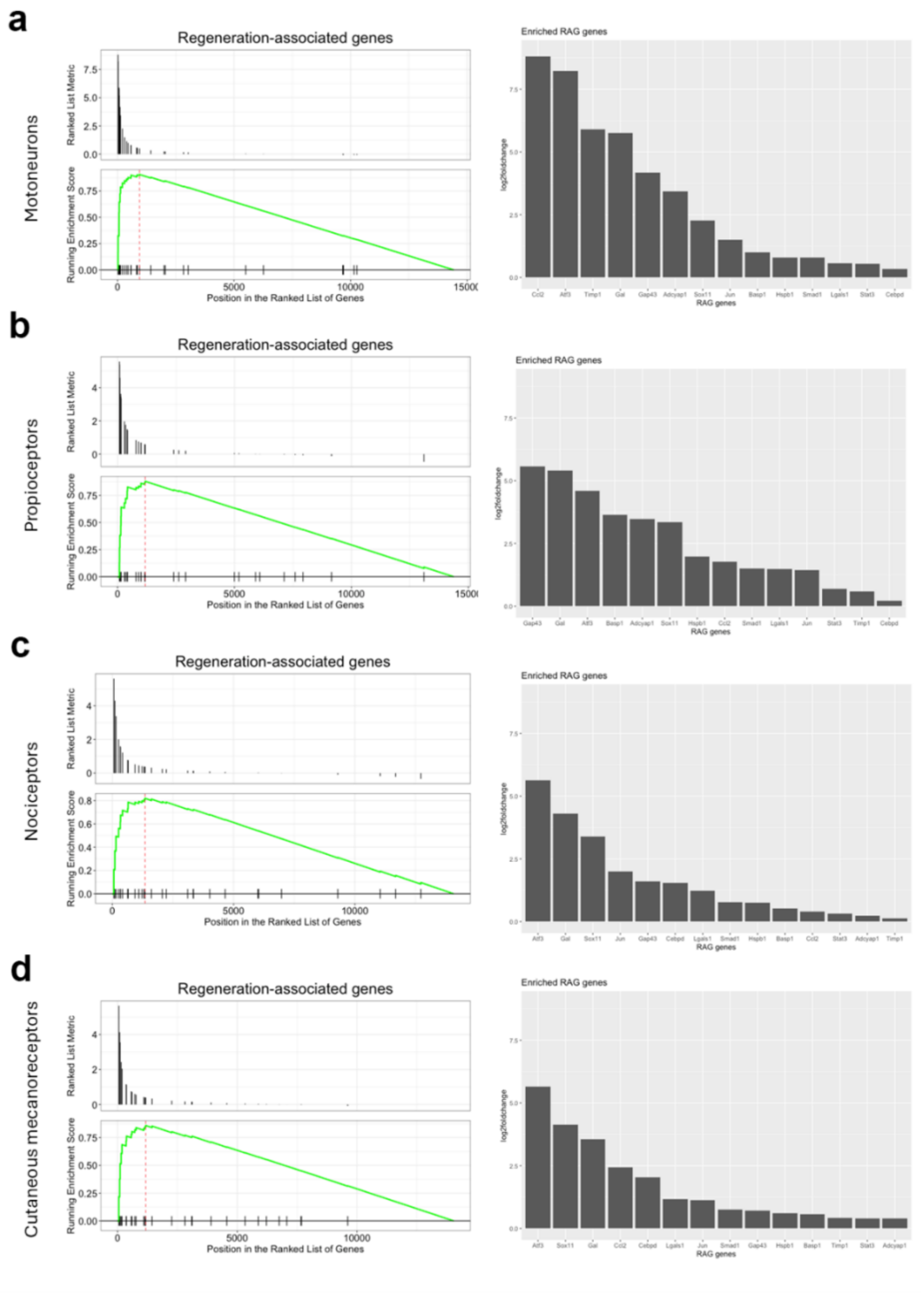


**Supplementary figure 2.** Gene set enrichment analysis of a literature-curated set of regeneration-associated genes in (a) motoneurons, (b) proprioceptors (Pvalb$^{+}$ neurons), (c) nociceptors (Trpv1$^{+}$ neurons), and (d) cutaneous mechanoreceptors (Npy2r$^{+}$ neurons).
